# Rhizome trait scaling relationships are modulated by growth conditions and are linked to plant fitness

**DOI:** 10.1101/2021.05.17.444226

**Authors:** Dinesh Thakur, Zuzana Münzbergová

## Abstract

**Background and Aim:** Rhizomes are important organs allowing many clonal plants to persist and reproduce under stressful climates with longer rhizomes indicating enhanced ability of the plants to spread vegetatively. We do not however, know either how rhizome construction costs change with increasing length or vary with environmental conditions.

**Methods:** We analysed the rhizome length vs mass scaling relationship, the plasticity in the scaling relationships, their genetic basis, and how scaling relationships are linked to plant fitness. We used data from 275 genotypes of a clonal grass *Festuca rubra* originating from 11 localities and cultivated under four contrasting climates. Data were analysed using standard major axis regression, mixed-effect regression models and a structural equation model.

**Key Results:** Rhizome construction costs increased (i.e., lower specific rhizome length) with increasing length. The trait scaling relationships were modulated by cultivation climate and its effects also interacted with the climate of origin of the experimental plants. With increasing length, increasing moisture led to a greater increase in rhizome construction costs. Plants with lower rhizome construction costs showed significantly higher fitness.

**Conclusions:** This study suggests that rhizome scaling relationships are plastic, but also show genetic differentiation and are linked to plant fitness. Therefore, to persist under variable environments modulation in scaling relationships could be an important plants’ strategy.

## INTRODUCTION

Plants optimise the allocation of limited resources to different plant parts to increase growth, survival and reproduction in a given environment (Reich et al., 1997; Wright et al., 2004; Díaz et al., 2016). Changes in environmental conditions may thus modify the allocation patterns via resource trade-offs, which, eventually, affect allocation to organs representing fitness, e.g., reproductive organs or total biomass (Springate and Kover, 2014; Halbritter et al., 2018). Elucidating the patterns of allocation and determining factors that cause changes in resource allocation patterns are, therefore, critical for understanding plant carbon economics and, hence, plant functioning and persistence (long term survival) under current and future climates.

To explore allocation strategies and developing plant economic spectra, trait scaling relationships are very important (Wright et al., 2004; Niklas et al., 2007; Enquist et al., 2015). In the last two decades, the use of these relationships in ecological research has greatly enhanced our understanding of how resources are allocated in different plant parts, such as, leaves, twigs or stems (Niklas et al., 2007; Fajardo, 2016; Husáková et al., 2018; Deng et al., 2020). The basic expectation behind the scaling relationships is that the various traits are not independent but, rather, are linked to other traits via trade-offs (Reich 2014). Based on the scaling relationships among various plant traits, a range of studies have demonstrated that allocation to some plant functions increases with increasing size (e.g., increased leaf construction cost for larger leaves (Niklas et al., 2007); increased investment into the stem relative to the leaves for larger trees (Poorter et al., 2015)). This indicates that plants allocate disproportionately more resources into specific plant functions with changes in size (Poorter et al., 2015).

Most studies on trait scaling relationships in plants deal with leaf traits (e.g., Reich et al., 1997, Wright et al., 2004, Niklas et al., 2007, Milla and Reich, 2007), with fewer studies dealing with stem traits (Poorter et al., 2015), root traits (Chen et al., 2019; Deng et al., 2020) and plant biomass fractions (above-ground and below-ground) (Niklas and Enquist, 2001; Husáková et al., 2018). There is, however, no such information for rhizomes and Goldberg et al., (2020) recently highlighted the strong need to understand the allocation of resources to clonal growth in plants.

Rhizomes are ecologically important plant organs responsible for vegetative spread in many plant species (Klimešová and De Bello, 2009; Duchoslavová and Jansa, 2018), with increased importance in stressful climates, for example, above the tree-line (Billings and Mooney, 1968; Körner, 2003). Rhizomes form bud banks (Ott et al., 2019) and enable species persistence by tolerating extreme conditions (e.g. very low or high temperatures or flooding) and promoting rapid growth during favourable seasons (since they also act as storage organs) (Billings and Mooney, 1968; Hull, 2008; Kinmonth-Schultz and Kim, 2011). Rhizomes also have important roles in processes such as below-ground carbon sequestration (De Deyn et al., 2008), community assembly (Weiher et al., 1998) and plant community resilience to disturbances (Hull, 2008; Speed et al., 2010).

The length of the rhizome per unit dry mass investment (i.e., specific rhizome length (SR_z_L = rhizome length / dry mass)) is a key trait indicative of costs of vegetative spread. Specific rhizome length was shown to vary with nutrient availability (Ye et al., 2006) and rhizome burial and breakage (Balestri and Lardicci 2014). Based on patterns reported in the literature on scaling relationships among above-ground traits (e.g., Reich et al., 1997, Wright et al., 2004, Niklas et al., 2007, Milla and Reich, 2007), the construction costs of rhizomes with increasing length can increase, decrease or be independent of their length. We are, however, not aware of any study distinguishing between these scenarios. Such knowledge is nonetheless important and has implications in understanding the ability of clonal plants to spread vegetatively and grow.

Climate is a major driver of plant functioning with vast literature on patterns, causes and consequences of functional trait variation along climatic gradients. By contrast, little is known about the effects of climate on trait scaling relationships (Vasseur et al., 2018). Specifically, earlier studies analysing variation in scaling relationships due to climatic factors (e.g., Xiang et al., 2013; Fajardo, 2016; Klimešová et al., 2017; Thakur et al., 2019) have not separated the effects of specific environmental factors on these relationships. This might be because the majority of the earlier studies was carried out in very complex natural environments where multiple factors (e.g., soil properties, climate, and genetic changes) act at the same time. We thus need experimental studies to disentangle the role of each factor separately.

In addition to allowing the separation of the effects of single environmental factors, the great advantage of experimental studies is that they allow assessment of the specific mechanisms determining the scaling relationships. Specifically, by studying plants from different original conditions cultivated under different environmental conditions, it is possible to assess the relative importance of genetic differentiation and phenotypic plasticity in determining the scaling relationships. Genetic differentiation has a role in the generation and maintenance of biodiversity and, thereby, insight into the evolutionary processes and the adaptive divergence of populations may be. gained (Weiher et al., 1998). On the other hand, phenotypic plasticity allows individual organisms to develop the appropriate traits that are better suited to the particular environment that they encounter (Yan et al., 2013). While studies separating these effects in terms of plant performance are common (e.g., (Münzbergová et al., 2017; Datta et al., 2017; Manzanedo et al., 2019), studies doing so in terms of trait scaling relationships are rare (but see Vasseur et al., 2018)).

Changes in trait values such as in specific root length (Kramer-Walter et al., 2016), specific leaf area (Liu et al., 2016) and rhizome density (Meyer and Schmid, 1999) have been shown to affect carbon allocation and thus be linked to whole plant fitness. Similarly, Vasseur et al., (2018) demonstrated that changes in trait scaling relationships may also lead to changes in carbon use and thus to changes in plant fitness. The effects of scaling relationships on fitness may also depend on environmental conditions determining the resource use efficiency of the plants (Anderson, 2016; Mota et al., 2018). Our knowledge on the effects of the scaling relationships on fitness and its changes within a changing environment is, however, still very limited.

To fill the above-mentioned gaps in our knowledge, we aim to understand rhizome length vs mass scaling relationships, in general, in order to test the effect of climate (the original and cultivation, representing genetic differentiation and phenotypic plasticity) on these relationships and explore how these relationships are related to plant fitness. Using *Festuca rubra* as a model plant, we addressed the following questions: (1) How do rhizome length and mass scale with each other? (2) Do trait relationships change with changes in temperature and moisture of growth conditions and plant origin? (3) Are rhizome trait scaling relationships linked to plant fitness?

We hypothesise that as with leaf trait scaling relationships, the rhizome length will fail to keep pace with increasing mass. We expect that the scaling relationships will vary with growth conditions, as a result of which the plants growing in warmer and wetter climates should have a lower rate of increase in tissue construction costs (because there is no thermal or water limitation). We also hypothesise that the trait relationships differ between plants from different original climates. Additionally, it is expected that lower rhizome construction cost in longer rhizomes would lead to higher plant fitness.

## MATERIALS AND METHODS

### Study species and system

The data used in this study are from Münzbergová et al., (2017) on a widespread clonal grass *Festuca rubra* L. It is distributed in temperate to tundra regions in both hemispheres as well as in tropical mountains (GBIF.org 2021, accessed on 07/03/2021). It reproduces vegetatively by forming both intra-vaginal and extra-vaginal tillers on rhizomes, but also reproduces by seeds (Münzbergová et al., 2017). While reproducing vegetatively, this species varies in the growth form and, consequently, resource capture strategies, creating a continuum of forms between ‘phalanx’ and ‘guerilla’ strategy (Skálová et al., 1997). Opposite to the ‘guerilla’ strategy, the ‘phalanx’ strategy is characterised by a decrease in the length of rhizome internodes and a greater rate of branching (Lopez et al., 1994). The ‘phalanx’ strategy is adopted in resource-rich environments while the ‘guerilla’ strategy is favoured in resource-poor conditions (Skálová et al., 1997).

The dataset used represented trait values of 275 genotypes (the genotype identification method is given in Šurinová et al,. 2019) collected from 11 localities (25 genotypes per locality). These localities are part of a climatic grid (the SeedClim Grid) of factorially-crossed temperature and precipitation gradients in western Norway (see details in Klanderud et al., 2015). All these genotypes were cultivated in four growth chambers (Votch, 2014), simulating the climatic conditions of extreme localities of the climate grid (cold/warm combined with dry/wet, (with circles in Figure 1) resulting in 11 localities × 25 genotypes × 4 climates, (i.e., 1100 individuals) were grown in the experiment. The conditions in the growth chambers mimic the spring to summer climate of each population in the field (the second half of April to the second half of June). Prior to cultivation, the plants were grown in a common climate for about seven months in total (four months in the garden and three months in the greenhouse) to remove most of the transgenerational effects (Münzbergová et al., 2017; see also Münzbergová et al., 2019).

**Figure 1:**
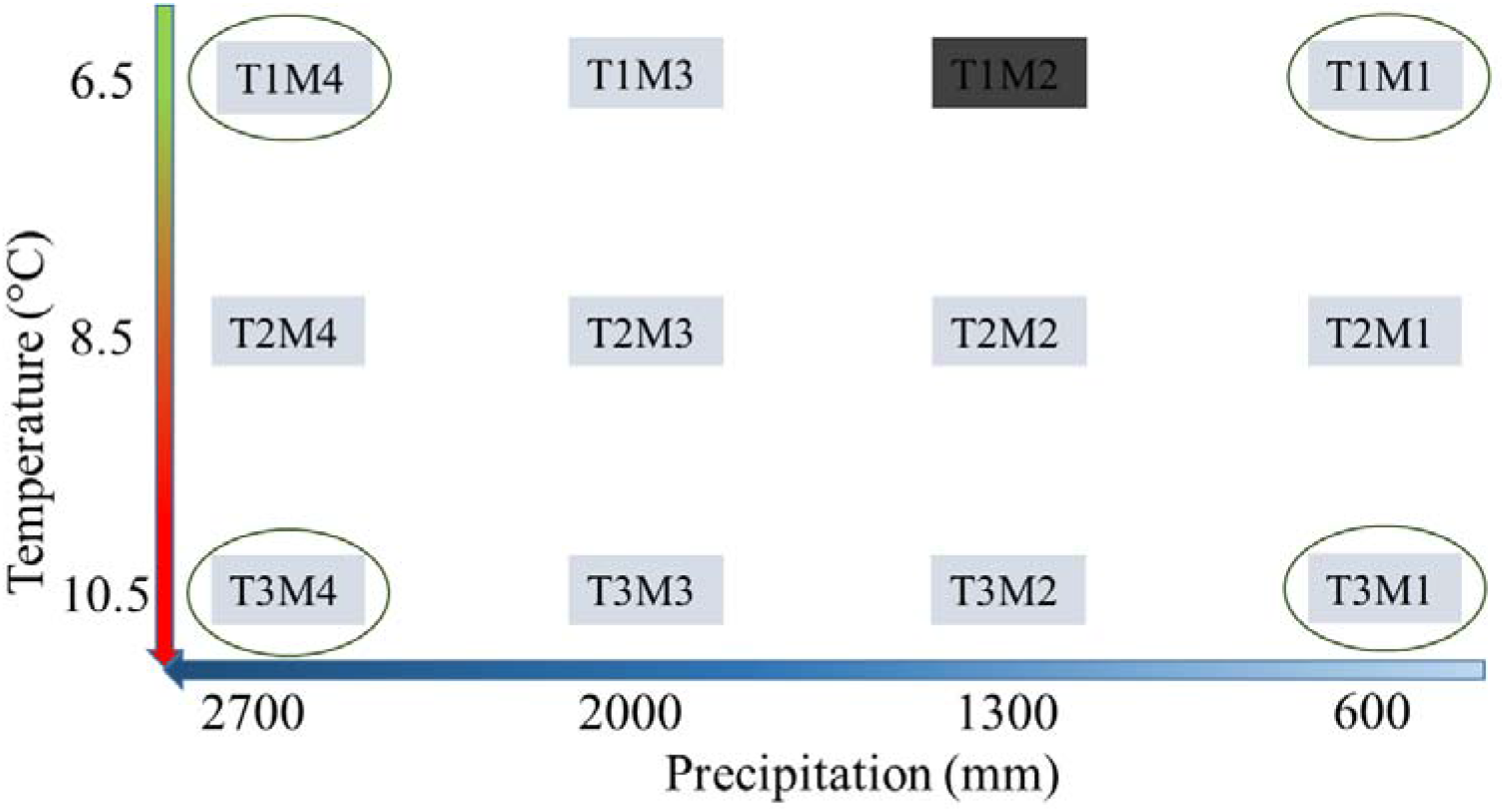
Representation of the climatic conditions of the localities from where the plant material was sampled. The four climates that are encircled were also used in climatic chambers as cultivation climates. The climatic condition in dark grey background (T1M2) was not used due to unavailability of the species. Temperature is mean for growing season and precipitation is annual.

### Growth conditions

The temperature in each the growth chambers followed the course of temperatures at the simulated natural locality (for details, see Münzbergová et al., 2017). The moisture level in the growth chambers was monitored as soil moisture using TMS dataloggers (Wild et al., 2019) (three inside each chamber) and then adding the necessary amount of water to mimic the soil moisture of the natural localities (also monitored using the TMS dataloggers). Each data-logger was placed in a pot with a growing *Festuca* plant identical to the experimental plants, but which was not a part of the experiment (not used for any other measurements). In the climate chambers with dry conditions, plants were watered with about 20 mL of tap water per plant, applied to the trays if the soil moisture was lower than 15%. In the wet regime, plants were cultivated under full soil saturation with ∼1.5 cm water level in the tray. By manipulating the soil moisture in the growth chambers, we are mimicking soil moisture at localities with a certain precipitation level. We thus refer to the moisture conditions in the growth chamber as precipitation values of the simulated localities. Day length and radiation levels were also controlled in a manner that these mimic the conditions of the original localities (more details in Münzbergová et al., 2017).

Some studies suggest that performing experiments in growth chambers lead to pseudo-replication of the experiment (Hurlbert, 1984). This argumentation has been later refuted as non-justified (e.g., Oksanen, 2001; Johnson et al., 2016). Most importantly, even Hurlbert (2004) stated that such a setting is not a problem in cases when the main focus is not on the main effects of the chamber climate, but on any interactive effects, such as the rhizome scaling relationships in our case (see (Münzbergová et al., 2017) for an extended discussion of the issue).

### Plant cultivation

The experiment ran from the end of February to the end of August 2015. The experiment was initiated by planting a single ramet without any rhizomes and with roots developed in water over three weeks prior to planting into each pot. Thanks to this, all below-ground structures of the experimental plants were of identical age.

In mid-June 2015, all the above-ground parts were cut to 3 cm to simulate mowing in natural conditions. After the final harvest in August 2015, we separated the plants into above-ground and below-ground parts; the below-ground parts were carefully sorted into roots and rhizomes. The total length of all rhizomes per plant was measured and all the biomass was then dried to a constant mass at 60 °C and weighed. The number of rhizomes per plant could not be counted due to their possible breaking during their extraction from the soil, so we only have information on their total length. This restricts us from mentioning if the total rhizome length is due to one rhizome or multiple rhizomes. This does not, however, have any influence on the results as, in any case, (i.e., the total length is due to one or more rhizomes) the overall longer length of the rhizome(s) per plant would reflect a greater ability to spread.

### Plant traits

This study primarily focused on analysing the relationship between rhizome length (mm) and rhizome mass (g). In addition, we also used other measured traits from the same individuals (i.e., above-ground biomass (g), number of ramets, total plant biomass (g), net photosynthetic rate (PN) and specific leaf area (SLA, mm^2^/mg) estimated by Münzbergová et al., (2017), Stojanova et al., (2018) and Kosová et al., (2021) to test if plant fitness and performance are linked to changes in the scaling relationships. Since this species reproduces predominantly clonally and only rarely flowered during the course of the study (Münzbergová et al., 2017), we used above-ground biomass, number of ramets and total plant biomass as estimates of plant fitness. Biomass is one of the widely used estimate of plant fitness (Younginger et al., 2017) and number of ramets is considered as good estimate of fitness in grasses (Pan and Price 2001). We also linked the scaling relationships to net photosynthetic rate (PN) and specific leaf area (SLA) to understand how changes in rhizome mass per unit length with increasing length is linked to changes in leaf ability to fix carbon and leaf construction cost. Due to the high workload required to measure these two traits, the data on these are only available from 10 genotypes originating from each of the four most extreme climates of the climatic grid: wettest and driest combined with warmest and coldest, i.e., 4 original climates × 4 cultivation climates × 10 genotypes = 160 measurements in total (the same as in Kosová et al., 2021).

### Data analyses

#### Standardised major axis regression

Standardised major axis (SMA) regression, a method commonly used for analysing scaling relationships, was used to calculate scaling exponent and elevation (α and β, respectively) for the relationship between rhizome length and mass (Niklas et al., 2007; Warton et al., 2012). SMA regression is most appropriate when both variables have errors and there is not a clear distinction between dependent and independent variables (Warton et al., 2012). A large number of earlier studies had used SMA to explore change in dry mass investment with increasing leaf area. It has been described using the formulae: *Area=* □β*Mass*^α^, where β is the elevation and α is the scaling exponent of the log-transformed area vs. mass regression curve (Niklas et al., 2007). In the case of rhizomes, the formulae will take the form *Length* □*=* □*β*_*1*_ *Mass*^α^. Since *SR*_*Z*_ *L* □*=* □*Length/Mass* and *Mass=* □*β*_*2*_ *Length*^α^, it follows that *SR*_*Z*_ *L=*□ *(1/β*_*2*_ *) Length*^*1-*α^. The value of α>1 in this equation would indicate that length fails to keep pace with mass (diminishing returns), whereas α<1 means the opposite (increasing returns).

Using the SMA regression, first, we estimated the scaling exponents and elevation for each cultivation climate across all populations in order to identify how the general scaling exponent is modulated by the cultivation climate. Second, we separately estimated the scaling exponents for plants from each original climate (population) grown at each cultivation climate to test if trait scaling relationships of plants from different original climates differ when grown in similar climate. The same analysis will also permit testing if the scaling exponents for plants from the same climate of origin differ when cultivated in different climates. In all the cases, we tested if the value of the scaling exponent is significantly different from 1 using one sample t-test. Using multiple post-hoc comparisons, we tested for differences in scaling exponents between plants grown in different cultivation climates and among plants from different original climates grown in different cultivation climates. SMA was done using ‘sma’ function with the argument ‘multcomp=TRUE’ (for multiple comparisons) in the ‘SMATR’ package (Warton et al., 2012) of R Version 4.0.3 (R Core Team and Core R Team, 2019).

Using SMA regression we also explored the effect of the direction of climate change on scaling relationships. For this, we estimated the change in temperature (ChangeT) and the change in moisture (ChangeM) by subtracting cultivation climate values from the original climate (expressed as °C for temperature and mm of rainfall, see above). For instance, if the original temperature is 6.5°C and the cultivation temperature is 12.5°C, the temperature change is +6°C (for more details see Münzbergová et al., 2017). We then estimated the scaling exponents based on differences in moisture or temperature between the original and the cultivation climate. Since scaling exponents differed significantly based on changes in the moisture, we also tested how change in moisture affects scaling exponents (direction of effect) using linear regression (‘lm’ function in R).

#### Mixed-effect models

While SMA regression is the best method to analyse the trait relationships, it does not allow study of the effects of different factors and their interactions in these relationships. Further, it does not allow consideration of any additional structure of the data (in our case, genotypes). We thus also analysed the data using mixed-effects regression models (see Husáková et al., 2018 for a similar approach). In the model, we used length as a dependent variable and mass as an explanatory variable, with original climate (temperature and precipitation, referred to as Otemp and Omois respectively), cultivation climate (referred to as Ttemp and Tmois) and interactions as fixed factors, and genotype as a random factor. Since the model was complex (with up to five interacting factors) we used the Akaike Information Criterion (AIC) to select the best model (Säfken et al,. 2018).

We also used the mixed effects regression to explore the effect of the direction of climate change on trait relationships, providing a more straightforward interpretation of the possible interactions between the climate of origin and that of cultivation. For this, we used ChangeT and ChangeM as fixed factors in the model and also tested their interactions with mass. As above, genotype was used as a random factor. Mixed-effects regression was carried n out using ‘lmer’ function in ‘lme4’ package (Bates et al., 2015) of R Version 4.0.3 (R Core Team and Core R Team 2019). Rhizome length and mass values were log-transformed before model fitting.

#### Relationships with fitness

To get an overview of the effect of the scaling exponent on plant fitness and resource capture traits, we used structural equation modelling (SEM) using the ‘sem’ function of the ‘lavaan’ package (Rosseel, 2012) of R. Data points in SEM were values of the scaling exponent, one for each population and cultivation conditions, and average trait values based on the same plants; one also for each population and cultivation conditions. In the SEM, we only used the variables that were significantly related to the scaling exponent. The relationship was tested using linear regression (‘lm’ function in R), (for details see supplementary information S1). We considered three possible measures of plant fitness in the SEM, the number of ramets and above-ground biomass and the total biomass. As the scaling exponent was not related to the number of ramets and above-ground biomass, only total biomass was used in the SEM presented. While doing the SEM, the best model was selected based on the lowest AIC value of the model and highest p value of Chi-Square (χ^2^) (Schermelleh-Engel et al., 2003; Barrett, 2007). The other measures of goodness of model fit (comparative fit index, Tucker Lewis index, root mean square error of approximation, and the standardised root mean square residual) as recommended by Coughlan et al., (2008) were also used to determine the model fit.

In SEM, we used the scaling exponent as the explanatory variable for response variables (rhizome length, SLA, and total biomass). We also used SLA as explanatory variable for PN and rhizome length. PN was also used as an explanatory variable for rhizome length. SLA and rhizome length were also used as explanatory variables for total biomass (indicator of fitness). Additionally, we also ran the model by using biomass excluding rhizomes and total biomass as indicators of fitness; the results were largely similar (not shown).

## RESULTS

### Effects of the cultivation climate on the scaling relationships

The mixed-effects regression results revealed that the relationship between rhizome length and mass was affected by both the moisture and the temperature of the cultivation climate (i.e., driven by phenotypic plasticity); the effect of moisture was, however, stronger (Table 1). The effects of temperature and moisture did not, however, interact (Table 1).

**Table 1.**
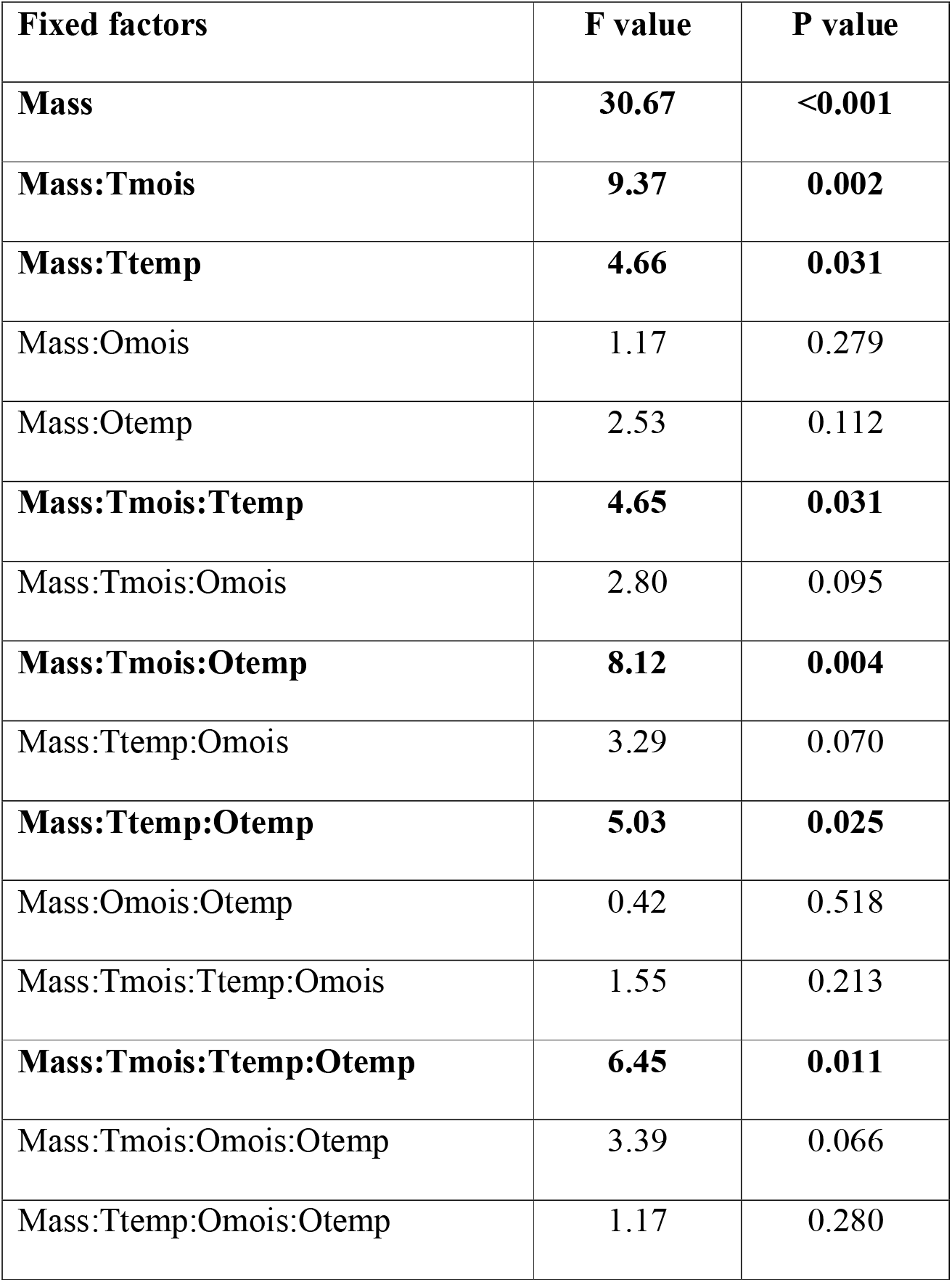
Linear mixed-effects regression model describing the effect of rhizome mass (g), original (O) and cultivation (T) climate (temperature and moisture) and their interactions on rhizome length (mm). Genotype was used as a random factor in the model. Only effects of mass alone and in interaction with the other factors are shown here. Full results of the model are given in supplementary Table S2.1. Significant values (p < 0.05) are shown in bold.

From the SMA regression, we found that plants cultivated in different cultivation climates differed significantly in the numerical values of α, see Table 2. The value of the scaling exponent (α) was significantly greater than 1 in all four cultivation climates (Table 2 and Figure 2). This indicates that rhizome dry mass increases disproportionately faster than rhizome length. In other words, rhizome construction costs increase with increasing length. As a consequence, longer rhizomes tend to have lower SR_z_L. The value of the scaling exponent (α) was significantly larger in plants cultivated in wet climates (both warm wet and cold wet) than in dry climates (Table 2 and Figure 2). This indicates that larger rhizomes are even more costly (in terms of biomass investment per unit length) in wet than in dry cultivation climates.

**Table 2.**
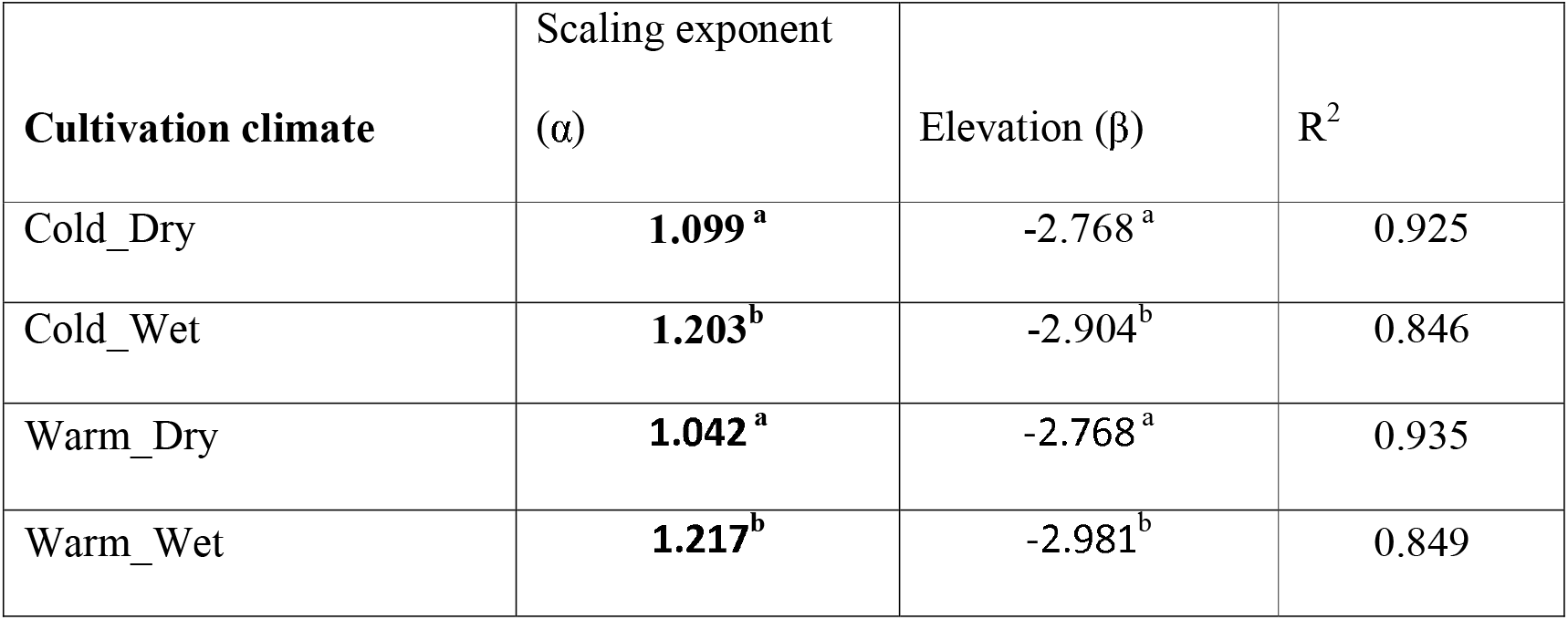
Scaling exponent (α) and elevation (β) for scaling relationship between rhizome length (mm) and rhizome mass (g) of *Festuca rubra*. The plants in different cultivation climates differ among each other in their scaling relationships. The value of scaling exponents in all the cases (presented in bold) were significantly greater than 1. 95% confidence intervals of α and β are given in supplementary Table S2.2. The superscript letters represent significant differences in scaling exponent among cultivation climates.

| Cultivation climate | Scaling exponent | | $R^2$ |
| --- | --- | --- | --- |
| | ($\alpha$) | Elevation ($\beta$) | |
| Cold_Dry | **1.099<sup>a</sup>** | -2.768<sup>a</sup> | 0.925 |
| Cold_Wet | **1.203<sup>b</sup>** | -2.904<sup>b</sup> | 0.846 |
| Warm_Dry | **1.042<sup>a</sup>** | -2.768<sup>a</sup> | 0.935 |
| Warm_Wet | **1.217<sup>b</sup>** | -2.981<sup>b</sup> | 0.849 |

**Figure 2:**
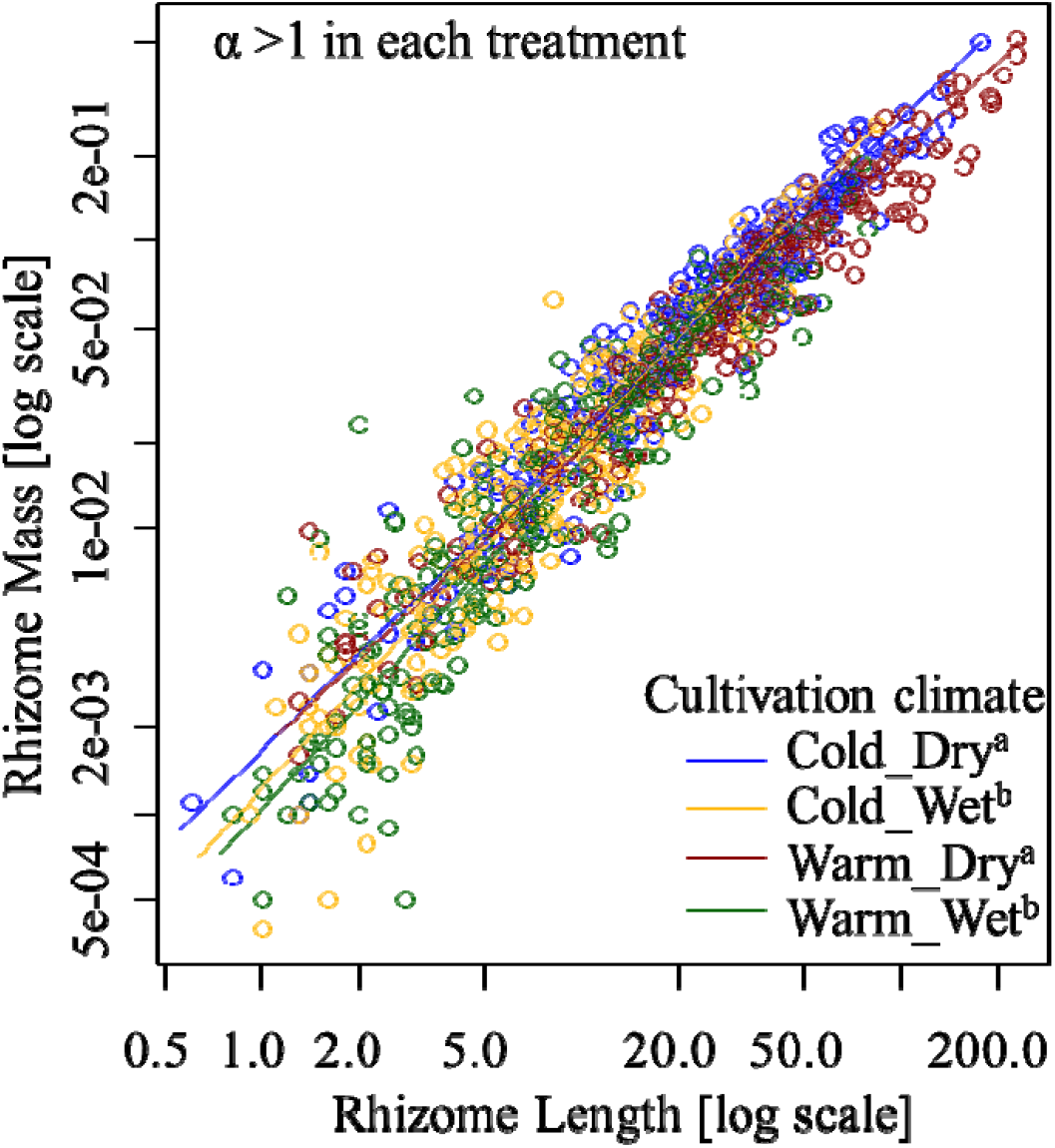
Relationship between rhizome length (mm) and mass (g) for plants from different cultivation climates. The superscript letters represent significant differences in scaling exponent among cultivation climates.

Despite significant differences in the scaling exponents among the four cultivation climates, the value of the scaling exponents was always significantly greater than 1 (Figure 2). This means that across various cultivation climates, the increases in rhizome mass result in disproportionately less gains in rhizome length (because α > 1,) but the gain is even lower in wet climates. When the value of the scaling exponents from each climate of origin in each cultivation climate were tested against the value of 1, it was found that the value of the scaling exponent was either equal to 1 or greater than 1 but was never less than 1. This indicated that there could be a proportionate increase in rhizome length and mass or there is a disproportionately higher increase in mass than length, but length can never increase proportionately greater than mass.

### Effects of the climate of origin on the scaling relationships

In the linear mixed-effects regression analysis, there were no significant interactions of mass with original climate only (Table 1). The only significant effects of the original climate were in interactions with the cultivation climate (i.e., Mass:Tmois:Otemp, Mass:Ttemp:Otemp, Mass:Tmois:Ttemp:Otemp in Table 1) indicating that plasticity in the relationships between rhizome length and rhizome mass depends upon the climate of origin. Additionally, in the SMA results, there were significant differences among plants from different original climates within a cultivation climate (evident only in T3_M1 cultivation climate -(see Table 3 and Figure 3) as well as between cultivation climates (e.g., T1_M1 and T3_M4 in Table 3). Scaling exponents also differed in plants from the same climate of origin when they were grown in different cultivation climates (e.g., in T2_M1 and T2_M4). The value of the scaling exponent for each original climate across all the cultivation climates ranged from 0.903 (statistically not different from 1; e.g., climate of origin T2_M4 grown in warm and dry climate, T3_M1) to 1.395 (significantly greater than 1 (e.g., genotypes from climate T3_M4 grown in cold and wet climate, T1_M4.

**Table 3.** Scaling exponent (α) and elevation (β) of scaling relationship between rhizome length (mm) and rhizome mass (g) based on the origin climate and the cultivation climate. In each case the value of R^2^ was > 0.647. The values of scaling exponents in bold are significantly greater than 1. 95% confidence intervals of α are given in supplementary Table S2.3. ** represents that scaling exponents differs among original climates for plants grown in the same cultivation climate (in the same column) and ^##^ represent that plants from the same original climate differ in scaling exponent among different cultivation climates (in the same row).

|  | Cultivation climate |  |  |  |  |  |  |  |
| --- | --- | --- | --- | --- | --- | --- | --- | --- |
|  | Cold_Dry<br>(T1_M1) |  | Cold_Wet<br>(T1_M4) |  | Warm_Dry<br>(T3_M1)** |  | Warm_Wet<br>(T3_M4) |  |
| Origin<br>climate | $\alpha$ | $\beta$ | $\alpha$ | $\beta$ | $\alpha$ | $\beta$ | $\alpha$ | $\beta$ |
| T1_M1 | 1.082 | -2.811 | **1.385** | -2.993 | **1.183** | -2.981 | **1.310** | -3.063 |
| T1_M3 | 0.997 | -2.620 | **1.201** | -2.906 | **1.156** | -2.980 | **1.265** | -2.994 |
| T1_M4 | 0.967 | -2.488 | **1.221** | -2.952 | 1.005 | -2.622 | 1.115 | -2.861 |
| T2_M1<sup>##</sup> | 1.111 | -2.771 | **1.253** | -2.884 | 0.982 | -2.652 | **1.281** | -2.986 |
| T2_M2 | 1.000 | -2.626 | 1.099 | -2.818 | **1.177** | -2.951 | 1.214 | -3.036 |
| T2_M3 | 1.088 | -2.793 | **1.287** | -2.988 | 1.034 | -2.774 | **1.324** | -3.072 |
| T2_M4<sup>##</sup> | **1.296** | -3.056 | **1.254** | -2.988 | 0.903 | -2.557 | **1.266** | -3.081 |
| T3_M1 | 1.101 | -2.833 | 1.083 | -2.785 | 0.999 | -2.733 | 1.006 | -2.734 |
| T3_M2 | 1.107 | -2.796 | 1.140 | -2.893 | 0.957 | -2.658 | 1.147 | -2.944 |
| T3_M3 | 1.085 | -2.754 | 1.153 | -2.814 | 1.085 | -2.799 | **1.199** | -3.090 |
| T3_M4 | 1.102 | -2.731 | **1.395** | -3.064 | 0.986 | -2.660 | 1.066 | -2.647 |

**Figure 3:**
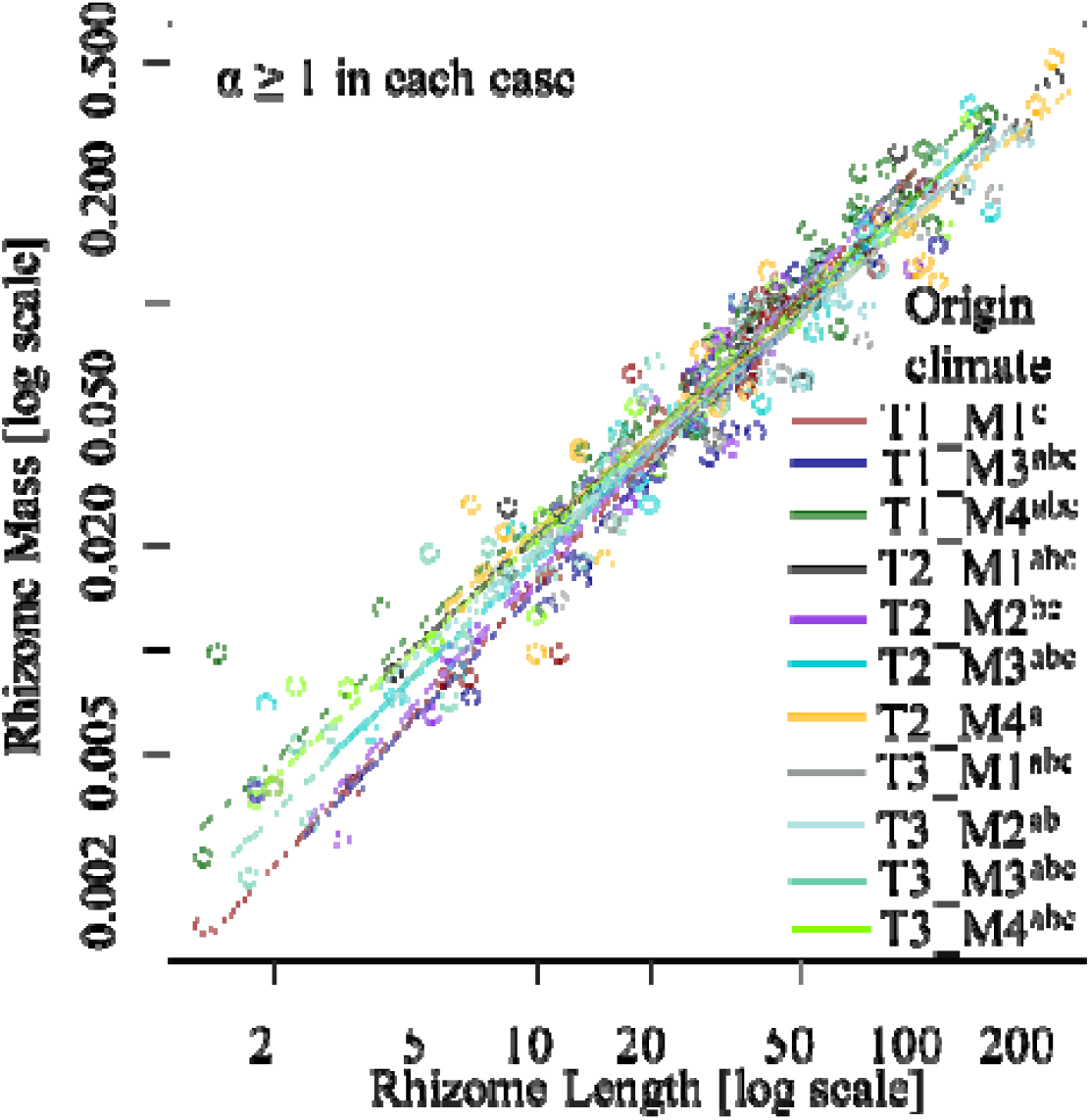
Differences in scaling exponent for the relationship between rhizome length (mm) and mass (g) among plants from different original climates grown under warm and dry cultivation climate. The plot is shown only for warm and dry cultivation climate because in other cultivation climates the scaling exponents were statistically not different among each other. T1 to T3 represent origin temperatures from low to high and M1 to M4 represent origin moistures from low to high (for details see Figure 1).

### Effects of climate change on scaling relationships

In the mixed-effects regression analysis based on degree of change in temperature and moisture between the climate of origin and the cultivation climate, significant interaction between ‘Mass’ and ‘ChangeM’ (change in moisture) was found (Table 4). This indicated differing scaling exponents based on changes in moisture. The SMA regression also revealed similar results and the value of the scaling exponent significantly increased with an increase in moisture (Figure 4). Change in temperature had no effect on the scaling exponent. This indicates that increasing or decreasing temperatures does not cause any additional change in rhizome construction cost with increasing length, but, with an increase in moisture, the rhizome mass (construction cost) increases disproportionately more than the rhizome length at an increased rate.

**Table 4:** Linear mixed-effects regression model statistics describing the effect of rhizome mass (g), climate change (temperature and moisture) and their interactions on rhizome length (mm). Genotype was used as a random factor. Only effects of mass alone and its interaction with the other factors are shown here. Full results of the model are given in supplementary Table 2.4. Significant values (p < 0.05) are shown in bold.

| Fixed factors | F value | P value |
| --- | --- | --- |
| Mass | **7853** | **< 0.001** |
| Mass:changeT | 2.40 | 0.136 |
| Mass:changeM | **20.51** | **<0.001** |
| Mass:changeT:changeM | 3.83 | 0.059 |

**Figure 4:**
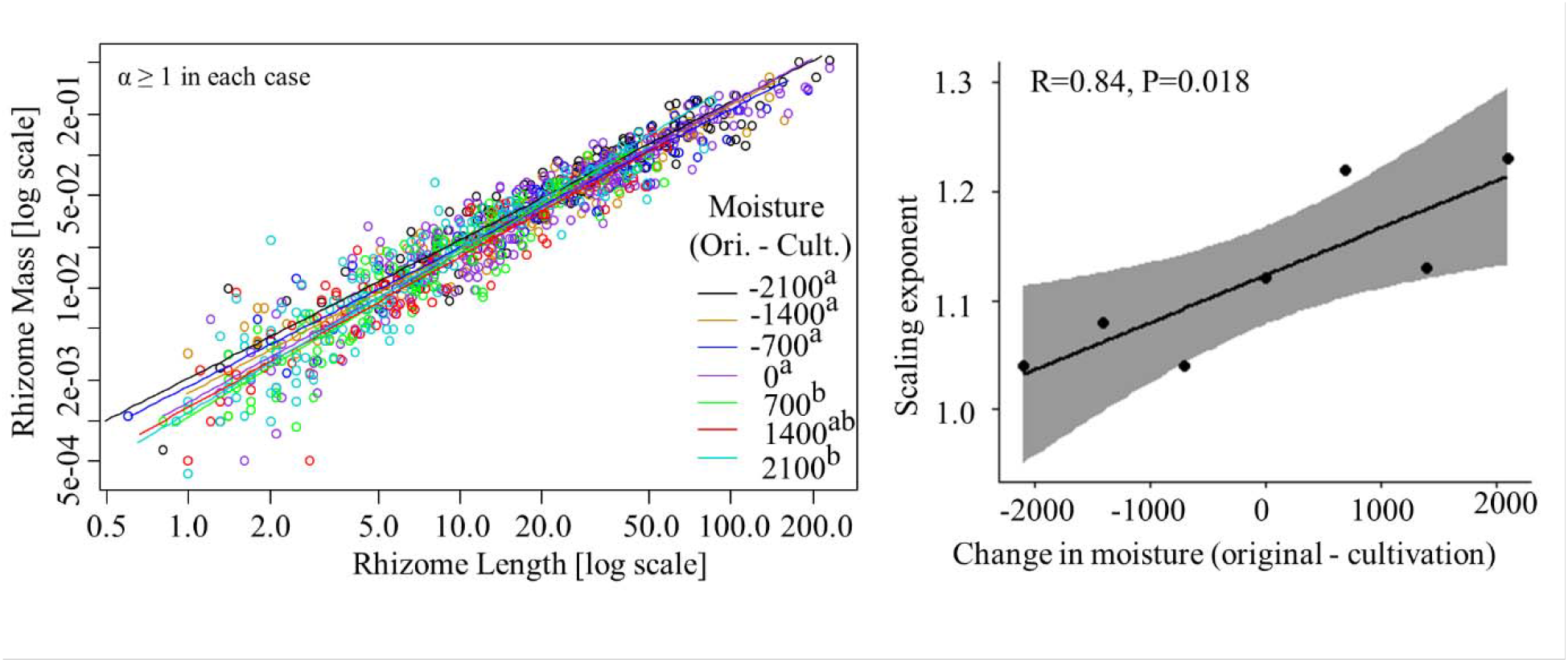
Figure in the left represents scaling relationship between rhizome length (mm) and mass (g) for plants cultivated under changed moisture (mm) conditions (differences in moisture between origin and cultivation climate). The superscript letters represent significant difference in scaling exponent. The right figure represents the change in scaling exponent with changes in moisture (original - cultivation).

### Relationship between scaling relationships and plant fitness

The high p value of the χ^2^ (0.99) as well as other estimated measures (e.g., Tucker Lewis index, root mean square error of approximation etc.) indicated a good fit of the structural equation model (supplementary Table S2.5). The model was able to moderately explain the variances in each of the response variables (R^2^ ranged from 0.34 to 0.68) (Figure 5). The model revealed that the increasing value of the scaling exponent significantly negatively influenced rhizome length and SLA, indicating that both rhizome length and SLA increases with decreasing value of the scaling exponent. The increasing SLA further led to an enhanced photosynthetic rate (PN) thereby increasing the total carbon fixed by the plant, but the PN was not significantly related to total biomass. When SLA were higher, total biomass also tends to be higher and was evident in the model from a marginally-significant, positive relationship (p=0.054) between the two. However, increasing SLA also led to decreasing rhizome length. Additionally, longer rhizomes were also significantly positively linked with higher plant total biomass.

**Figure 5:**
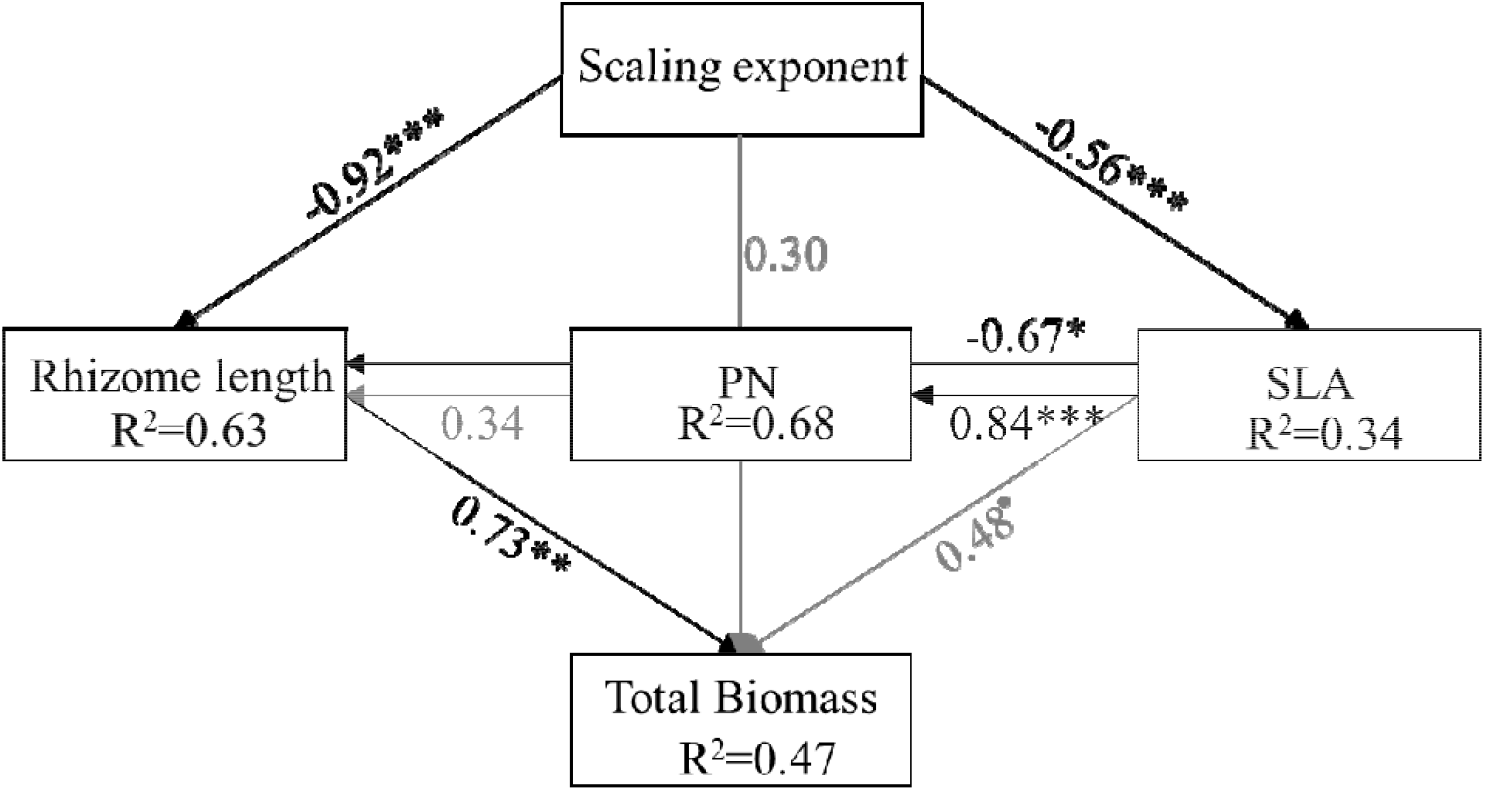
Structural equation model representing the effect of scaling exponent on traits related to plant fitness. PN = net photosynthetic rate, SLA = Specific leaf area (mm^2^/mg). Path coefficients between variables are unstandardized regression coefficients and part of the variances explained by the model (R^2^) are given under the variable names. The black arrows with regression coefficients in black letters are significant regressions (p ≤ 0.05), while those in grey are non-significant (p > 0.05). *** indicates p <0.001, * indicates p<0.05 and • indicates p<0.1. Goodness of fit statistics of the model are given in supplementary Table S2.5.

## DISCUSSION

Trait scaling relationships have important implications for understanding the plant resource use strategies and the ability of plants to acclimatise or adapt to variable environmental conditions. In this study, we analysed the rhizome length versus mass scaling relationships and found a considerable variation in the scaling exponents. The main findings emanating from this study are: (i) The rhizome length fails to keep pace with increasing mass and, as a result, with increasing rhizome length the rhizome construction costs increase. (ii) Trait scaling relationships are modulated by climatic conditions in which the plants grow, and their effect interacts with the climate of origin. (iii) Climate change affects the scaling relationships and wetter climates lead to greater increase in rhizome construction costs with increasing length. (iv) Trait scaling relationships are closely linked to plant performance (e.g., carbon capture capacity by leaves).

### Rhizome construction cost increases with increasing length

Our analyses show that scaling relationships for the functional traits that primarily influence plant vegetative spread have a scaling exponent greater than one in all the cultivation climates. Thus, changes in rhizome length fail to keep pace with increasing mass. This indicates that either bulk rhizome-tissue density or thickness (or both) increase as rhizome length increases (causing decreased SR_Z_L). The result is similar to those reported for above-ground plant traits (e.g., leaf size and leaf mass, Leishman et al., 2007; Price and Enquist, 2007; Atkin et al., 2008)). Theoretically, these results mean diminishing returns from an increase in rhizome length (i.e., increasing construction cost for larger rhizomes). This finding is in parallel with Niklas et al., (2007) and (Milla and Reich 2007) who reported ‘diminishing returns’ in the case of leaves.

The finding follows the metabolic scaling theory (Enquist et al., 2007) supporting the fact that trait relationships are similar across evolutionary-distinct organisms as well as in ecologically distinct functional traits. We assume that this increasing cost could be because rhizomes are also the organs to transport water and minerals to above-ground parts, additional mechanical support is, therefore, required for longer distance transport. Additionally, rhizomes are also storage organs in grasses (for soluble as well as for non-soluble carbohydrates (Klimeš et al., 1999; Kinmonth-Schultz and Kim, 2011), but this feature is unlikely to cause higher mass per unit length with increasing length. Although such information is missing in the literature, we also do not have any reason to expect that storage function of rhizomes would affect scaling between the rhizome mass and the length (i.e., larger rhizomes should store disproportionately more or less carbohydrates per unit length than smaller ones). Nevertheless, testing if rhizome length and stored carbohydrates have isometric relationship is an open question.

Further, the study species is commonly grazed in natural habitats, so longer rhizomes are more vulnerable to herbivore damage via trampling. Therefore, greater investment in mechanical tissues (more xylem or thick-walled cells) may also be helpful in this case. In this context, Striker et al., (2006) stated that the porosity of roots in a grass species (*Paspalum dilatatum*) is lower (density is higher) when exposed to trampling. Although there is additional cost associated with longer rhizomes, longer rhizomes enable positioning of new ramets away from the parents thereby decreasing intra-clone competition and increasing resource availability and the higher investment is thus reasonable. This might be an important strategy of clonal plants and may lead to overall positive returns (e.g., higher dominance in a community).

### Scaling relationships are modulated by growth conditions

Rhizome length vs mass scaling relationships varied among plants cultivated under different climatic conditions demonstrating that scaling relationships are modulated in response to growth conditions. While there are no previous studies on rhizome scaling relationships allowing comparison with our results, several previous studies showed the effect of climate on scaling relationships of leaf traits (e.g., Wright et al., 2005; Atkin et al., 2008; Xiang et al., 2013; Thakur et al., 2019). All these studies were, however, from natural environments and were not able to separate phenotypic plasticity from genetic differentiation. By using data from experimentally-manipulated conditions, we were able to separate multiple effects in this study. Our results provide strong evidence that plant scaling relationships are plastic. This finding is important because evidence of the plasticity in scaling exponents is lacking, particularly in plants (for animals, see Casasa & Moczek, 2019)). Specifically, these findings indicate the importance of modulation in scaling relationships under variable climates in *Festuca rubra*. We argue that similar to the role of trait plasticity in plant persistence under variable climates, plasticity in scaling relationships is also important and might be helpful in optimal resource allocation under changing climatic conditions to enhance plant fitness.

The variation in scaling relationships with changing cultivation conditions might help the plant to balance the need for efficient vegetative spread, resource use and protection against desiccation and soil herbivores. The direction of the shift in the scaling exponent (lower value in drier sites) was, however, unexpected. The majority of the previous literature indicates that warmer and wetter conditions favour plant growth (Natali et al., 2012; Buermann et al., 2018) but our data indicate the opposite (because higher tissue construction costs have negative consequences). This could be because the wet climate was too wet (simulation of 2700 mm of annual rainfall) and thus not favourable for the species as also claimed by Münzbergová et al., (2017). These too wet conditions may create hypoxic conditions; it is known that plants in hypoxic conditions have thick, aerenchymatic below-ground structures (Pedersen et al., 2021). To increase thickness there would be additional allocation of resources in the aerenchyma (which otherwise is not needed) and when rhizomes are longer, more thicker tissues can be helpful for longer-distance, oxygen transport. Therefore, more carbon allocation is needed with increasing rhizome length. Secondly, a transition towards a ‘phalanx’ resource use strategy (higher number of ramets per unit length) was also evident in this species in wet climates (Supplementary Figure S2.1) and this transition could be responsible for the higher rate of increase in rhizome construction cost with increasing length. This can be expected because plants with ‘phalanx’ strategy have shorter internodes, which are linked to increased stem density, and decreased hydraulic conductivity (Jacobsen et al., 2020). The decrease in hydraulic conductivity with increasing number of nodes is because some of the vessels end in each internode thus disrupting the resource flow (Jacobsen et al., 2020). Therefore, with increasing rhizome length, the need to maintain hydraulic conductivity of plants in wet conditions having many internodes may impose additional costs. Third, with increasing wetness, there might be increased activity of soil herbivores or parasitic fungi (Velásquez et al., 2018). Hence, additional investments per unit length are needed for protection because tissues with lower carbon content are easier to degrade (Silver and Miya, 2001). Overall, our results indicate that it is more economical for this clonal species to form longer rhizomes in conditions with lower water availability. This was evident in the results as plants in drier conditions had longer rhizomes (Supplementary Figure S2.2).

The differences in the value of scaling exponents among cultivation climates were apparently very small but, as shown by Milla & Reich (2007), the small differences in the numerical value of the scaling exponent (among cultivation climates in our case) can translate into very large differences in construction cost per unit length when rhizomes differ greatly in length. These small differences can have huge consequences for the overall plant performance (e.g., vegetative spread) because of greater cost to build and maintain a unit of rhizome length, which should constrain the maximum rhizome length achieved by a plant. Milla and Reich (2007) demonstrated that 1.07 value of the scaling exponent in leaf area vs mass relationship cause 22% increase in per unit leaf construction cost, when leaf size increased by about 6.5-fold (i.e., 96 to 622 cm^2^). Considering this example, one can imagine the impact of only a small deviation of the scaling exponent from 1.

Interestingly, in this study, the total length of the rhizome in climates with a greater increase in construction cost (e.g., warm and wet) remained far less than in climates with a lesser increase in construction cost (see Supplementary Figure S2.2). We argue that there might be economic constraints that limit rhizome elongation (or ability to spread) when construction cost is increasing rapidly. This shorter length of the rhizome in warm and wet climates may, therefore, be the negative consequence of greater increase in construction cost.

Our result also hints that plants tend to keep the rhizome construction cost under a certain threshold, above which rhizomes are not formed. This was supported by the fact that SR_z_L was not significantly lower in warm and wet cultivation climate than dry and wet climate, but rhizome length was kept short. Moreover, despite shorter rhizomes, the SR_z_L at the cultivation climates with higher construction costs have higher variance (inferred based on trait driver theory (Enquist et al., 2015), (Supplementary Figure S2.3). This indicated that in such climates, plants are beginning to construct rhizomes with a lower construction cost but, as a consequence of a larger value of the scaling exponent, construction cost surpassed to that in drier climates at much shorter lengths.

### Genetic differentiation in the scaling relationship

We suspected that scaling relationships might also be result of adaptation to environments at the places of origin. While we did not find any significant interaction between mass and the climate of origin, we detected triple interactions between mass, the climate of origin and that of cultivation indicating genetic differentiation in the plasticity of the scaling relationships. This finding provided supporting evidence to possible local adaptation in scaling relationships associated with phenotypic optimisation to enhance fitness. This is in line with Vasseur et al., (2018) providing evidence for a genetic basis in the scaling relationship of plant dry mass vs growth rate in *Arabidopsis thaliana*.

In combination with the effect of climate change on the scaling relationships, our results support the idea that scaling relationships are modulated significantly by original as well as cultivation climate at intraspecific levels. Therefore, we suspect that, at community level, differences in scaling relationships (reported in many earlier studies on leaf traits (Niklas and Enquist, 2001; Milla and Reich, 2007; Niklas et al., 2007; Thakur et al., 2019)) could be result of both intraspecific (i.e., responses to climate and local adaptation) as well as interspecific effects (i.e., due to species turnover).

### Scaling relationships are linked to plant fitness

The inverse relationship of the scaling exponent with the total plant biomass indicated that plant fitness decreases when formation of longer rhizomes becomes more costly (i.e., a larger value of the scaling exponent). This was also expected because higher investment per unit length can have negative consequences (less investment) for other traits involved in resource acquisition. The SEM result supports that decreased scaling exponent was associated with lower leaf construction cost (i.e. SLA). In turn, this lower leaf construction cost was associated with higher photosynthetic efficiency. Therefore, when leaf construction costs are lower, more light can be intercepted by the same aboveground mass (Milla and Reich 2007) and, subsequently, more carbon is fixed. Overall, performance of leaves was greatly increased when the scaling exponent was lower.

Our results also indicated that when the value of scaling exponent is lower, longer rhizomes are formed indicating the plant’s ability to vegetatively spread is also enhanced. Additionally, the net returns as represented by total biomass were also dependent on rhizome length (the greater the length, the higher the returns), (see also supplementary Figure S1.1). The negative relationship between the SLA and rhizome length in the SEM indicates that the ‘exploitative’ resource use strategy of leaves is linked to the limited ability of plants to spread vegetatively. The probable reason is that when leaves are more productive, more resources are allocated to roots to exploit more nutrients from the soil. This was supported by a strong positive relationship between SLA and root biomass (Supplementary figure S2.4).

Our results on the relationship of the scaling exponent with leaf traits (PN and SLA) also indicate that changes in below-ground tissue construction costs are linked to above-ground tissue construction costs. We could have tested this hypothesis in this study, but the available data of leaf area and leaf mass did not allow us to do so (as the SLA was based on data from leaf fragments). Overall, the findings of this study provided evidence that when scaling relationships are modulated, plant fitness is compromised. The scaling exponents indicating ‘diminishing returns’ (as defined in Niklas et al., 2007) had negative consequences on plant fitness. As we have very scarce information about plant investments into rhizomes (in comparison with other plant organs) and how these investments are modulated by growth conditions, more such studies are awaited to know if the patterns are similar across clonal species with different phylogenetic histories. While many current studies are dealing with the benefits of clonal growth, more studies (such as this one) are needed to shed light on clonal growth costs. It can be expected that as with above-ground plant organs, the patterns observed in this study may also stand true for other species that reproduce mainly clonally. Nevertheless, this needs to be tested in diverse species occurring in different ecoregions.

## Conclusions

Our analysis of the scaling of rhizome length vs mass indicates that rhizome length fails to scale one to one with rhizome mass due to which longer rhizomes have higher construction costs (i.e., lower SR_z_L). Our findings provided evidence for plasticity in scaling relationships in plant traits and demonstrated that the rate of increase in rhizome construction costs with length depends upon the growth conditions. Among the climatic conditions, the moisture of the cultivation conditions was the main determinant of the scaling relationships. Our results also provided additional support for genetic basis in the scaling relationship. Finally, we have demonstrated that scaling relationships are closely linked to plant performance and lower scaling exponents (with length, a lower increase in construction cost) are linked to traits representing acquisitive resource use strategy.

## Supporting information

Complete supplementary information

## ACKNOWLEDGEMENTS

We thank the POPEKOL discussion group for useful comments on the manuscript. The study was supported partly by the project GACR 19-00522S and partly by institutional research projects RVO 67985939 and MSMT. We also thank Vigdis Vandvik and her team for granting access to the SEEDCLIM climate grid from where the initial material had been obtained and Věroslava Hadincová for her help with the collection of the initial data.

