## Supplementary material for "Rhizome trait scaling relationships are modulated by growth conditions and are linked to plant fitness": Complete supplementary information

**Supplementary information S1**

***Methods: Relationships between scaling exponent and traits***

To test for relationship between traits and scaling exponent, we used linear regression (using ‘lm()’ function in R) with aboveground biomass, number of ramets, PN, SLA, root biomass and total plant biomass as dependent variable and scaling exponent as explanatory variable. PN was log transformed to achieve normality prior to regression analysis. An increase in the value of these traits with decreasing scaling exponent would indicate that lower value of scaling exponent lead to higher plant fitness or performance and *vice-versa.*

In the same way, we also tested the relationship between rhizome length and rhizome mass and scaling exponent. In this analysis, a change in length with value of scaling exponent would indicates that length is not constant but is modulated depending upon construction cost. Similarly, change in mass with scaling exponent would indicate that density or thickness or length or all three are modulated. An inverse relationship between the length and scaling exponent in this case would mean that when construction cost is lower rhizomes are longer (more ability to spread vegetatively). But, if at the same time, mass is also inversely related with scaling exponent, it means that plant is also investing more in rhizomes to make very long rhizomes (even more ability to spread) which are also more dense or thicker.

***Results: Relationship between scaling relationships and plant traits***

In the linear model regression, it was found that neither the aboveground biomass nor the number of ramets was affected by value of scaling exponent. However, total plant biomass (representative of total carbon captured) was negatively affected by the value of scaling exponent indicating that with the increase in rhizome construction cost the plant is able to fix less amount of carbon (Figure S1.1). Similarly, root biomass also decreased with increasing value of scaling exponent indicating that the plants ability to acquire nutrients (represented by root biomass) increased with decreasing scaling exponent. It was intriguing, as to how plant fixed more carbon without no extra investment in aboveground tissues. So, when we did linear regression, it was revealed that PN and SLA were negatively related with scaling exponent (Figure S1.1). The relationship with SLA indicated that with decreasing rhizome construction cost leaf construction cost also decreased (i.e. plant can form more leaf in same amount of biomass). In the same way, relationship with PN indicated increasing photosynthetic efficiency with decreasing construction cost. With increasing value of scaling exponent, both length and mass of rhizome decreased (Figure S1.2) indicating that plants invested more in rhizomes and ability to vegetative spread (increase in rhizome length) increases with decrease in scaling exponent.

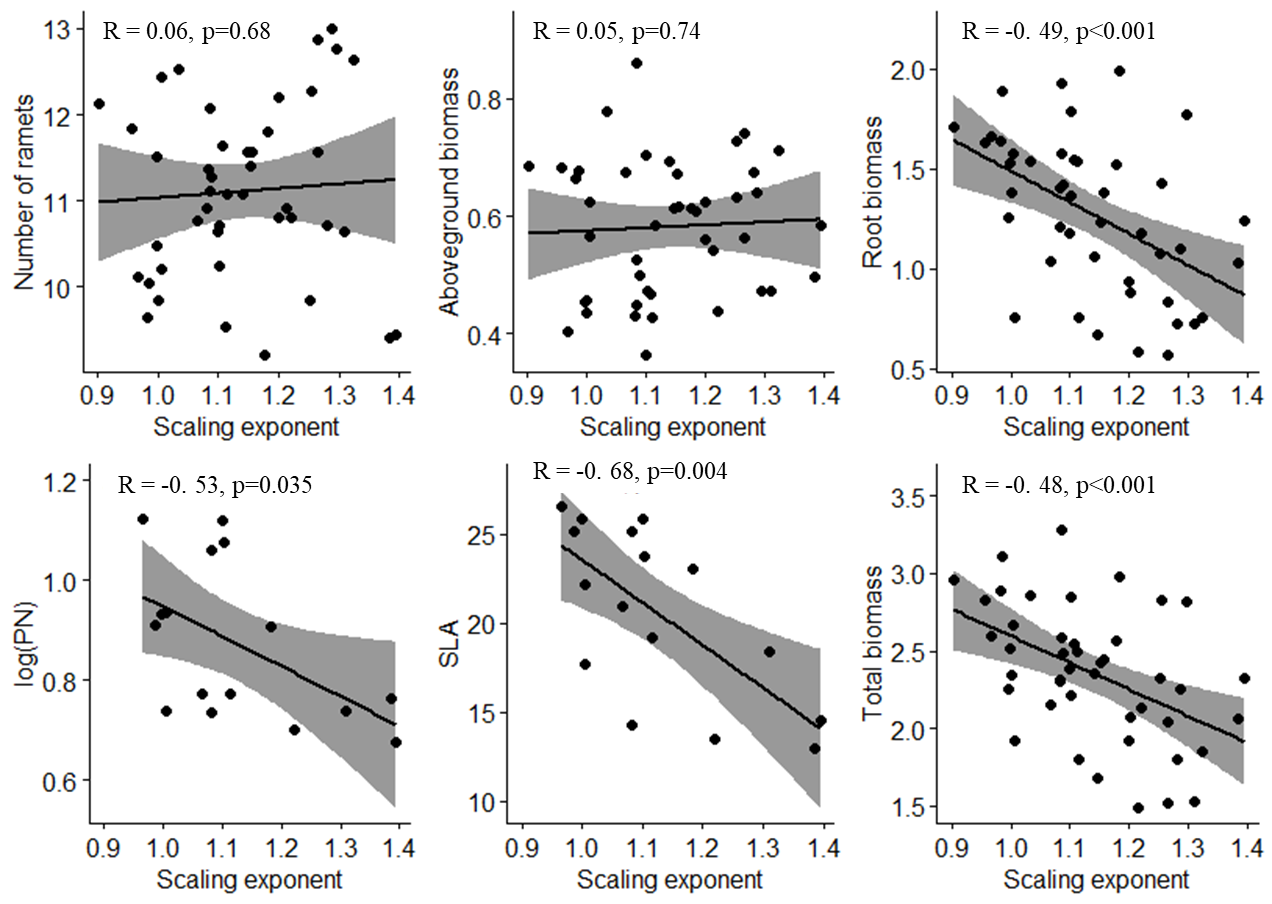

**Figure S1.1:** Regression plots showing the dependence of plant traits on scaling exponent.

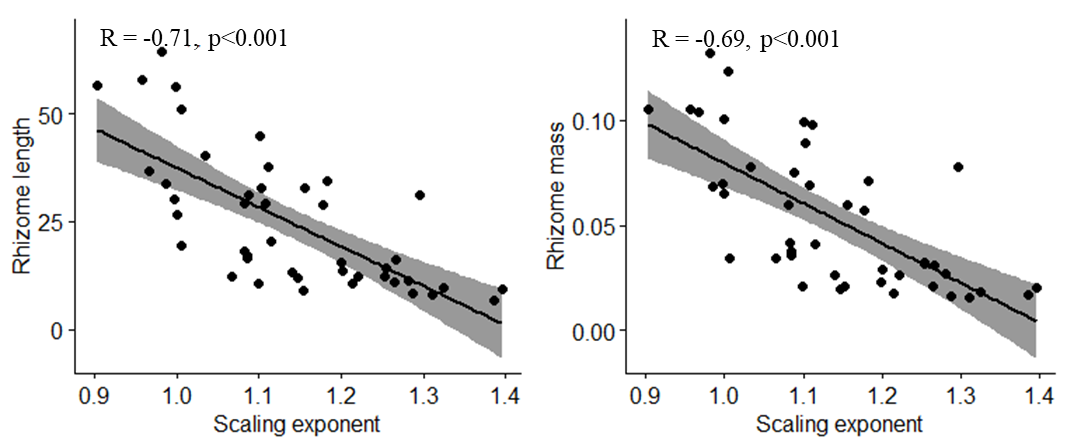

**Figure S1.2:** Regression plots showing the dependence of rhizome length and mass traits on scaling exponent.

**Supplementary Information S2**

**Table S2.1**: Linear mixed effects regression model describing the effect of rhizome mass, original (O) and cultivation (T) climate (temperature and moisture) and their interactions on rhizome length. Genotype was used as a random factor in the model.

| **Fixed factors** | **F value** | **P value** |
| --- | --- | --- |
| Mass | 30.67 | <0.001 |
| Tmois | 13.95 | <0.001 |
| Ttemp | 3.77 | 0.052 |
| Omois | 3.48 | 0.063 |
| Otemp | 4.12 | 0.043 |
| Mass:Tmois | 9.37 | 0.002 |
| Mass:Ttemp | 4.66 | 0.031 |
| Mass:Omois | 1.17 | 0.279 |
| Mass:Otemp | 2.53 | 0.112 |
| Tmois:Ttemp | 3.95 | 0.047 |
| Tmois:Omois | 7.89 | 0.005 |
| Tmois:Otemp | 12.06 | 0.001 |
| Ttemp:Omois | 4.07 | 0.044 |
| Ttemp:Otemp | 5.83 | 0.016 |
| Omois:Otemp | 1.73 | 0.189 |
| Mass:Tmois:Ttemp | 4.65 | 0.031 |
| Mass:Tmois:Omois | 2.80 | 0.095 |
| Mass:Tmois:Otemp | 8.12 | 0.004 |
| Mass:Ttemp:Omois | 3.29 | 0.070 |
| Mass:Ttemp:Otemp | 5.03 | 0.025 |
| Mass:Omois:Otemp | 0.42 | 0.518 |
| Tmois:Ttemp:Omois | 1.81 | 0.179 |
| Tmois:Ttemp:Otemp | 5.45 | 0.020 |
| Tmois:Omois:Otemp | 8.05 | 0.005 |
| Ttemp:Omois:Otemp | 1.80 | 0.180 |
| Mass:Tmois:Ttemp:Omois | 1.55 | 0.213 |
| Mass:Tmois:Ttemp:Otemp | 6.45 | 0.011 |
| Mass:Tmois:Omois:Otemp | 3.39 | 0.066 |
| Mass:Ttemp:Omois:Otemp | 1.17 | 0.280 |
| Tmois:Ttemp:Omois:Otemp | 0.58 | 0.447 |

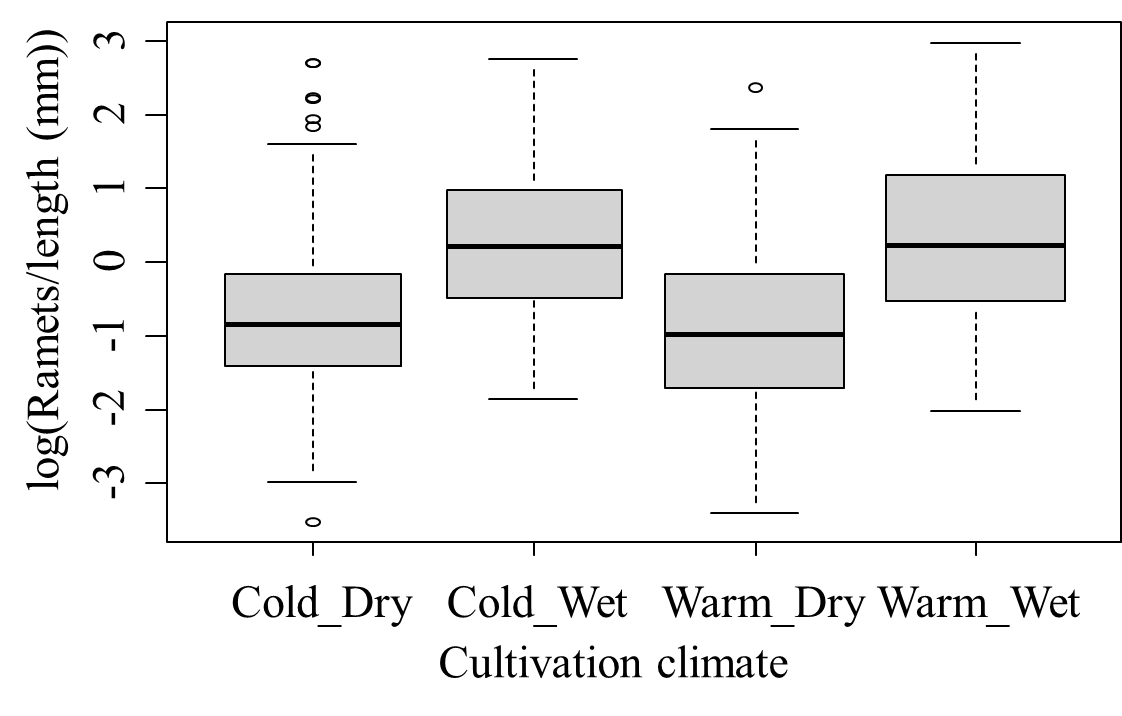

**Figure S2.1:** Boxplots showing the distribution of ramets per unit rhizome length in plants cultivated at four different cultivation climates. The higher values in this plot indicate shift towards 'phalanx' resource use strategy (i.e. longer internodes and greater rate of branching). Therefore, plants in wet climates are more towards 'phalanx' resource use strategy.

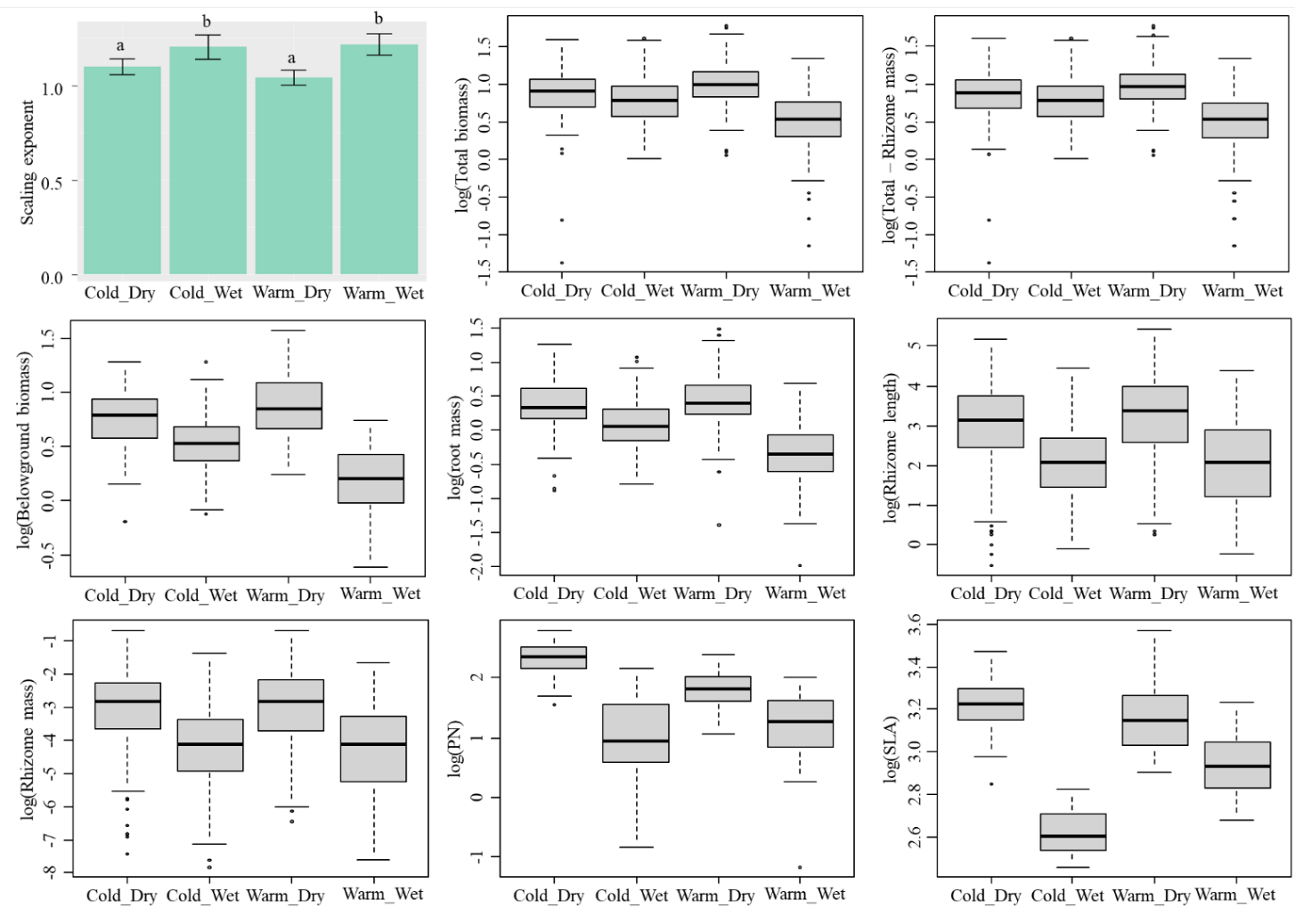

**Figure S2.2:** Representation of the value of scaling exponent for four different cultivation climates (uppermost left) and boxplots showing the distribution of various traits at these cultivation climates.

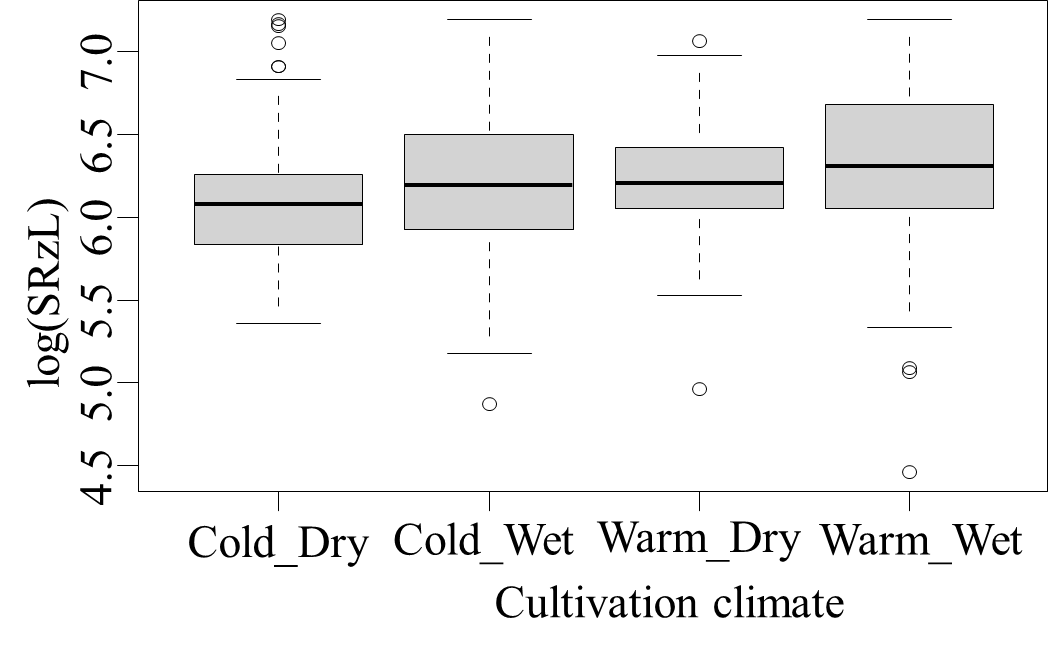

**Figure S2.3:** Boxplots representing distribution of specific rhizome length for different cultivation climates. **Table S2.2.** Elevation (β) and scaling exponent (α) for scaling relationship between rhizome length and rhizome mass of *Festuca rubra*.

| **Cultivation climate** | **β** | **95% CI of β** | **α** | **95% CI of α** | **R^2^** |
| --- | --- | --- | --- | --- | --- |
| Cold_Dry | -2.768 | -2.826 to -2.709 | 1.099 | 1.058 to 1.141 | 0.925 |
| Cold_Wet | -2.904 | -2.966 to -2.842 | 1.203 | 1.143 to 1.267 | 0.846 |
| Warm_Dry | -2.768 | -2.825 to -2.710 | 1.042 | 1.004 to 1.081 | 0.935 |
| Warm_Wet | -2.981 | -3.039 to -2.922 | 1.217 | 1.162 to 1.274 | 0.849 |

**Table S2.3.** Scaling exponent (α) and elevation (β) of scaling relationship between rhizome length and rhizome mass based on origin climate and cultivation climate. 95% confidence interval (CI) of both exponent and elevation are presented below the estimated value. In each case the value of R^2^ was more than 0.647.

| **Cultivation climate →** |  | **Cold_Dry (T1_M1)** | **Cold_Wet (T1_M4)** | **Warm_Dry (T3_M1)** | **Warm_Wet (T3_M4)** |
| --- | --- | --- | --- | --- | --- |
| **↓Origin climate** | | | | | |
|  |  | Estimate and  95% CI | Estimate and  95% CI | Estimate and  95% CI | Estimate and  95% CI |
| **T1_M1** | **α** | 1.082  0.944 to 1.239 | 1.385  1.180 to 1.625 | 1.183  1.081 to 1.296 | 1.310  1.119 to 1.533 |
|  | **β** | -2.811  -3.013 to -2.608 | -2.993  -3.176 to -2.810 | -2.981  -3.139 to -2.823 | -3.063  -3.224 to -2.903 |
| **T1_M3** | **α** | 0.997  0.842 to 1.181 | 1.201  1.034 to 1.394 | 1.156  1.021 to 1.308 | 1.265  1.059 to 1.513 |
|  | **β** | -2.620  -2.851 to -2.388 | -2.906  -3.100 to -2.711 | -2.980  -3.185 to -2.776 | -2.994  -3.213 to -2.775 |
| **T1_M4** | **α** | 0.967  0.821 to 1.140 | 1.221  1.045 to 1.427 | 1.005  0.887 to 1.139 | 1.115  0.932 to 1.333 |
|  | **β** | -2.488  -2.729 to -2.247 | -2.952  -3.156 to -2.748 | -2.622  -2.828 to -2.416 | -2.861  -3.109 to -2.612 |
| **T2_M1** | **α** | 1.111  0.971 to 1.272 | 1.253  1.017 to 1.545 | 0.982  0.887 to 1.086 | 1.281  1.117 to 1.469 |
|  | **β** | -2.772  -2.997 to -2.546 | -2.884  -3.151 to -2.618 | -2.652  -2.819 to -2.485 | -2.986  -3.159 to -2.812 |
| **T2_M2** | **α** | 1.000  0.897 to 1.115 | 1.099  0.955 to 1.266 | 1.177  1.056 to 1.313 | 1.214  0.972 to 1.516 |
|  | **β** | -2.626  -2.769 to -2.484 | -2.818  -2.968 to -2.669 | -2.951  -3.123 to -2.779 | -3.036  -3.324 to -2.749 |
| **T2_M3** | **α** | 1.088  0.981 to 1.206 | 1.287  1.086 to 1.525 | 1.034  0.924 to 1.156 | 1.324  1.153 to1.520 |
|  | **β** | -2.753  -2.916 to -2.590 | -2.988  -3.178 to -2.797 | -2.772  -2.940 to -2.603 | -3.072  -3.246 to -2.898 |
| **T2_M4** | **α** | 1.296  1.139 to 1.475 | 1.254  1.057 to 1.487 | 0.903  0.779 to 1.047 | 1.266  1.097 to 1.462 |
|  | **β** | -3.056  -30287 to -2.825 | -2.988  -3.224 to -2.752 | -2.557  -2.769 to -2.344 | -3.081  -3.281 to -2.881 |
| **T3_M1** | **α** | 1.101  0.943 to 1.286 | 1.083  0.919 to 1.277 | 0.999  0.881 to 1.132 | 1.006  0.862 to 1.176 |
|  | **β** | -2.833  -3.104 to -2.563 | -2.785  -2.989 to -2.582 | -2.733  -2.939 to -2.527 | -2.734  -2.919 to -2.550 |
| **T3_M2** | **α** | 1.107  0.991 to 1.236 | 1.140  0.924 to 1.406 | 0.957  0.851 to 1.077 | 1.147  0.990 to 1.331 |
|  | **β** | -2.796  -2.970 to -2.622 | -2.894  -3.136 to -2.651 | -2.658  -2.850 to -2.465 | -2.944  -3.103 to -2.784 |
| **T3_M3** | **α** | 1.085  0.946 to 1.245 | 1.153  0.966 to 1.375 | 1.085  0.975 to 1.206 | 1.199  1.051 to 1.367 |
|  | **β** | -2.754  -2.922 to -2.986 | -2.814  -2.993 to -2.634 | -2.799  -2.936 to -2.663 | -3.090  -3.267 to -2.914 |
| **T3_M4** | **α** | 1.102  0.958 to 1.267 | 1.395  1.031 to 1.888 | 0.986  0.914 to 1.063 | 1.066  0.812 to 1.399 |
|  | **β** | -2.731  -2.949 to -2.513 | -3.064  -3.477 to -2.652 | -2.660  -2.762 to -2.558 | -2.647  -2.963 to -2.331 |

**Table S2.4:** Linear mixed effects regression model statistics describing the effect of rhizome mass, climate change (temperature (changeT) and moisture (changeM)) and their interactions on rhizome length. Genotype was used as a random factor.

| **Fixed effects:** | **F value** | **P value** |
| --- | --- | --- |
| Mass | 7853 | < 0.001 |
| changeT | 8.29 | 0.004 |
| changeM | 18.26 | < 0.001 |
| Mass:changeT | 2.40 | 0.121 |
| Mass:changeM | 20.51 | < 0.001 |
| changeT:changeM | 2.83 | 0.093 |
| Mass:changeT:changeM | 3.83 | 0.051 |

**Table S2.5:** The values of estimated parameters indicative of goodness of fit of the model against the recommended cut-off for a good fit (Coughlan et al. 2008). All the measures indicated good fit of the model.

| **Measure of model fit** | **Value in current model** | **Cut-off for good fit** |
| --- | --- | --- |
| P value of Model Chi-Square (χ2) | 0.93 | p-value > 0.05 |
| Comparative Fit Index (CFI) | 1 | ≥ 0.90 |
| Tucker Lewis index (TLI) | 1.133 | ≥ 0.95 |
| Root Mean Square Error of Approximation (RMSEA) | <0.001 | < 0.08 |
| (Standardized) Root Mean Square Residual (SRMR) | 0.006 | < 0.08 |

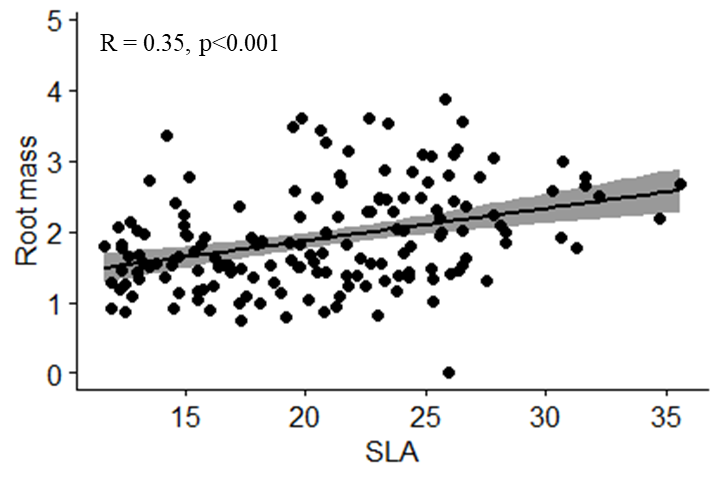

**Figure S2.2.4**: Scatterplot representing relationship between SLA (mm^2^/mg) and root mass (g).
